# Not one hormone or another: Aggression differentially affects progesterone and testosterone in a South American ovenbird

**DOI:** 10.1101/371690

**Authors:** Nicolas M. Adreani, Wolfgang Goymann, Lucia Mentesana

**Affiliations:** Department of Behavioural Neurobiology, Max Planck Institute for Ornithology, Seewiesen, Germany; Research Group Evolutionary Physiology, Max Planck Institute for Ornithology, Seewiesen, Germany

## Abstract

Behaviors such as territorial interactions among individuals can modulate vertebrate physiology and *vice versa*. Testosterone has been pointed out as a key hormone that can be rapidly affected by aggressive interactions. However, experimental evidence for such a link is mixed. In addition, behaviors can elicit changes in multiple hormones, which in turn have the potential to synergistically feedback to behavior. For example testosterone and progesterone can act interdependently in modulating male behavior. However, if aggression can affect progesterone levels in males remain unknown and – to the best of our knowledge – no one has yet tackled if and how aggressive behavior simultaneously affects testosterone and progesterone in free-living animals. We addressed these questions by performing simulated territorial intrusion experiments measuring both hormones and their ratio in male rufous horneros (Aves, *Furnarius rufus)* during the mating and parental care periods. Aggression affected testosterone and progesterone differentially depending on the period of testing: challenged birds had higher levels of progesterone during the mating period and lower levels of testosterone during parental care compared to controls. Challenged individuals had similar progesterone to testosterone ratios during both periods and these ratios were higher than those of control birds. In summary, territorial aggression triggered hormonal pathways differentially depending on the stage of the breeding cycle, but equally altered their ratio independent of it. Our results indicate that multiple related hormones could be playing a role rather than each hormone alone in response to social interactions.

## Introduction

Hormones can modulate behavior, but also behavior can feed back into physiology and generate a hormonal response of the organism, priming the organism for future behavioral interactions. In addition, a specific behavior can simultaneously elicit changes in multiple hormones (e.g.Elekonich and Wingfield, 2000). Given the pleiotropic nature of hormones, that is their capacity to affect multiple processes that range from development (reviewed by Harris 2015) to neural modulation (Harris-Warrick and Marder, 1991), and the fact that different hormones can interact with each other (Crews and Moore, 2005; Erickson et al., 1967; Godwin and Crews, 1997), it is relevant to gain insights regarding the effect that a specific behavior has on different hormones.

The relationship between aggressive behavior and testosterone levels has been widely studied in the field of behavioral endocrinology (reviewed by Hirschenhauser and Oliveira, 2006 and Oliveira, 2004), with the ‘challenge hypothesis’ providing a well-defined framework for grasping how social interactions can affect testosterone responses in a wide range of animals (e.g.Wingfield *et al.* 1990, Oliveira 2004). One of the core assumptions of the ‘challenge hypothesis’ is that testosterone mediates the trade-off between male-male competition and male contributions to parental care. As a consequence, males of biparental species should have a large androgen responsiveness, that is, they should maintain low breeding baseline levels of testosterone, unless they get involved in a male-male challenge. When this is the case, testosterone should rise to maximal levels and return to baseline after the interaction has ended, particularly so in the parental phase (Wingfield et al., 1990). Since its initial proposition ‘the challenge hypothesis’ has been modified to account for life-history variation across species (e.g. Wingfield & Hunt 2002; Goymann, Landys & Wingfield 2007; Lynn 2008), but its basic predictions for bi-parental species remained unchanged. Support for the challenge hypothesis has been found in different vertebrate taxa (reviewed by Hirschenhauser and Oliveira, 2006 and Oliveira, 2004). However, for birds - possibly the most intensively studied group - there is surprisingly little experimental support for the ‘challenge hypothesis’ and a strong bias towards northern hemisphere species (reviewed by Goymann, 2009 and Goymann et al., 2015).

Aggressive behavior has also been proposed to affect progesterone, another steroid hormone typically related to reproductive behavior in vertebrates (Beach, 1976; Komisaruk, 1967; Lindzey and Crews, 1986; Moore, 1978). Despite being a hormone associated mainly with female physiology, progesterone seems to be positively related with male aggressive interactions in vertebrates (Fuxjager et al., 2009; Schneider et al., 2003; Weiss and Moore, 2004; but see Erpino and Chappelle, 1971; Fraile et al., 1988, 1987). While for females there is a well-defined framework that links aggression with progesterone (‘progesterone challenge’ hypothesis) and proposes a suppressive effect of progesterone (Davis and Marler, 2003), if and how aggression affects progesterone in males is almost completely unknown.

Progesterone can also interact with testosterone in modulating male behavior (Crews et al., 1996; Erickson et al., 1967; Erpino, 1969; Erpino and Chappelle, 1971). For example, in male ring doves (*Streptopelia capicola*) progesterone inhibits sexual behavior when administered in combination with testosterone (Erickson et al., 1967). In contrast, in male whiptail lizards (*Cnemidophorus inornatus*) progesterone acts synergistically with testosterone to modulate sexual behavior (Crews et al., 1996). Remarkably, despite the potential interaction between both hormones the effect of male aggressive behavior on both hormones has not been investigated yet.

Rufous horneros (*Furnarius rufus*, hereafter termed hornero) are birds that inhabit the temperate zone of eastern South America and aggressively defend their territories year-round (Fraga, 1980). They are socially monogamous seasonal breeders and both sexes cooperate in nest-building, incubation and feeding the young (Fraga, 1980; Massoni et al., 2012). By comparing hormone levels of freely behaving male horneros that were challenged by a simulated intruder with control birds during two stages of the breeding cycle (mating and parental care periods), we tested whether territorial interactions would affect testosterone and progesterone levels, and the ratio between these hormones.

Based on the ‘challenge hypothesis’, we predicted that STIs would generate an increase in testosterone levels, in particular during the parenting period. Because the effect of aggression on progesterone levels has been experimentally studied only in females (Davis and Marler, 2003; Elekonich and Wingfield, 2000; Goymann et al., 2008), we based our predictions from the ‘progesterone challenge’ hypothesis on data from females. We thus expected that STIs would elicit a decrease in progesterone. Finally, if the predictions for testosterone and progesterone hold true, the ratio between progesterone and testosterone should decrease.

## Materials and methods

### (I) Experimental design

Fieldwork was carried out on the campus of INIA Las Brujas (National Institute of Agricultural Research), Canelones, Uruguay (34°40’ S, 56°20’ W; 0-35 m a.s.l.). We conducted the simulated territorial intrusions (STI) in two periods during the year 2016: during August-September when birds were building nests (“mating period”), and during November-December when pairs were feeding nestlings (“parental care period”).

We performed the STIs using one male stuffed hornero while playing back male solo songs and duets extracted from the database xeno-canto in ‘wav’ format (www.xeno-canto.org) during 20 minutes. From a pool of ten male solo-songs, ten duets and ten ‘pause’ files of 7-15 seconds duration, a new random choice of sounds was produced every time for each control and STI. In this way, the acoustic component of the STI differed for each territory, while the visual decoy stimulus was the same for each pair for ethical reasons. In other birds such as great tits (*Parus major*), mounted decoys (N = 15) did not explain variance in male aggressive behavior (Araya-Ajoy and Dingemanse, 2013) and since horneros are monomorphic in body size and plumage coloration (Diniz et al., 2016) it is unlikely that the usage of only one decoy influenced our results.

Because the contributions to the duet are sex-specific (Laje and Mindlin, 2003; Roper, 2005) observers could assign the sex to each focal individual. To validate this, molecular sexing was performed for every bird captured (following Griffiths et al., 1998). Observers assigned the sexes correctly in 26 out of 27 cases where they could assign a sex to the bird that was in the net. In every territory, during 20 min of STI each bird of the pair was observed focally by one of the two observers. Focal birds were assigned randomly to each observer (male or female). The traits recorded were: ‘latency of first approach’, ‘number of duets’, ‘number of solo song’, ‘time within 5 meters of the decoy’, ‘time on the nest’ and ‘number of flights over the decoy’. In a multivariate analysis of these characters as part of a different study, ‘flights over decoy’ and ‘number of duets’ were the most reliable predictors of the multivariate trait ‘aggression’ in horneros (Adreani et al., unpublished). Given that these two variables are highly correlated we related hormone concentrations to ‘flights over decoy’ only. After 20 minutes, mist nets were opened and the STI continued until the birds were captured (mean capture time ± sd = 5.94 ± 4.68 min after opening the nets). Control birds were caught with the same setup, however, in this case the mist nets were opened before the playback started and birds were caught before ten minutes of playback had passed (mean capture time ± sd = 4.50 ± 3.75 min after onset of playback). During the entire experiment we did not have repeated measures across controls and STIs, or across mating and parental phase.

Immediately after capture, for all birds, we took a blood sample (mean handling time ± sd = 4.75 ± 1.48 min) from the wing vein to determine post-capture testosterone and progesterone concentrations. Then, we injected 50 μl saline containing 5 μg of gonadotropin-releasing hormone (GnRH; Bachem, H3106) into the breast muscle. The GnRH injection served as a positive control to determine if birds would have been physiologically capable to increase testosterone concentrations in case they did not increase testosterone during the simulated territorial intrusion (for details see Goymann 2009 and Apfelbeck & Goymann 2011). The concentration of the GnRH injection was based on the amount used in previous studies (Apfelbeck and Goymann, 2011; Jawor et al., 2006) and adjusted according to the average body mass of horneros (~60 g). Tarsus length, wing length and weight were measured, and 30 minutes after GnRH injection a second blood sample was taken to measure GnRH-induced hormone concentrations, before releasing the birds.

Blood samples were centrifuged at 3000 rpm for 10 min. Plasma was extracted and promptly frozen at −20°C. After three weeks samples were transferred to a −150°C freezer where they were kept for six months and finally transferred to a −80°C freezer where they stayed for five weeks before analysis. Overall, we collected data from 24 control and 22 STI males sampled during the mating period; and 7 control and 9 STI males sampled during parental care. In addition, STI and control groups did not differ in body condition (Table S1), capture time (time since the end of the 20’ STI until the capture for the STI birds and time since start of the playback until capture for the controls - Table S2) and sampling time (time since the bird was trapped into the mist net until blood sample was taken - Table S2).

### (II) Hormone analysis

Each plasma sample was divided into two aliquots for testosterone and progesterone quantification respectively. Both hormones were extracted from the plasma and concentrations were determined by radioimmunoassay following the procedures described by Goymann et al. (2008). Mean recovery during extraction of the samples was 87.00% (+/- 0.19) for testosterone and 84.00% (+/- 0.05) for progesterone. One assay was run per hormone with lower detection limits of 3.5 pg/ml for testosterone and 22.1 pg/ml for progesterone. All samples used in our analyses were above these detection limits. The intra-assay variation was calculated from of an extracted chicken pool and was 3.5% for testosterone and 11.1% for progesterone.

### (III) Statistical analyses

The analyses were performed in R (R Core Team, 2013) using the statistical packages ‘*lme4*’ (Bates et al., 2014) and ‘*arm*’ (Gelman, Andrew; Yu-Sung, 2015) in a Bayesian framework with non-informative priors. We assumed a Gaussian error distribution, which was confirmed for all response variables after visual inspection of model residuals. We subsequently used the ‘*sim*’ function to simulate values from the posterior distributions of model parameters. We extracted the 95% credible interval (CrI) around the mean (Gelman and Hill, 2007), representing the uncertainty around our estimates. In all figures we display raw data, and the predicted estimate from the models and 2.5% - 97.5% CrI. We considered an effect to be statistically meaningful when the posterior probability of the mean difference between compared estimates was higher than 0.95 or when the estimated CrI did not include zero (for further details on statistical inference see Korner-Nievergelt et al., 2015). From the figures, a statistically meaningful difference between groups can be assumed if the CrI of one group do not overlap with the mean estimate of the other.

#### Body condition, capture time and handling time control

We performed a linear model for each body condition and capture time parameters (i.e., wing and tarsus length, body mass, capture and handling time). The explanatory variables in all models were: treatment (control vs. STI) and period (mating vs. parental care period). For details on these models see Supplementary Table 1 and Supplementary Table 2.

#### Effect of STI on hormones

We performed a linear mixed effect model for each hormone and the ratio between progesterone and testosterone. The three dependent variables were log-transformed. The explanatory variables in all models were: treatment (control vs. STI), period (mating vs. parental care period) and moment of sampling (*pre* vs. *post* GnRH injection). Finally, as each bird was blood sampled twice, ‘ID’ was included as random factor. For details on these models see Table 1. We initially considered the effect of handling time, but because of its negligible effect it was excluded from the final models to avoid overfitting.

**Table 1.**
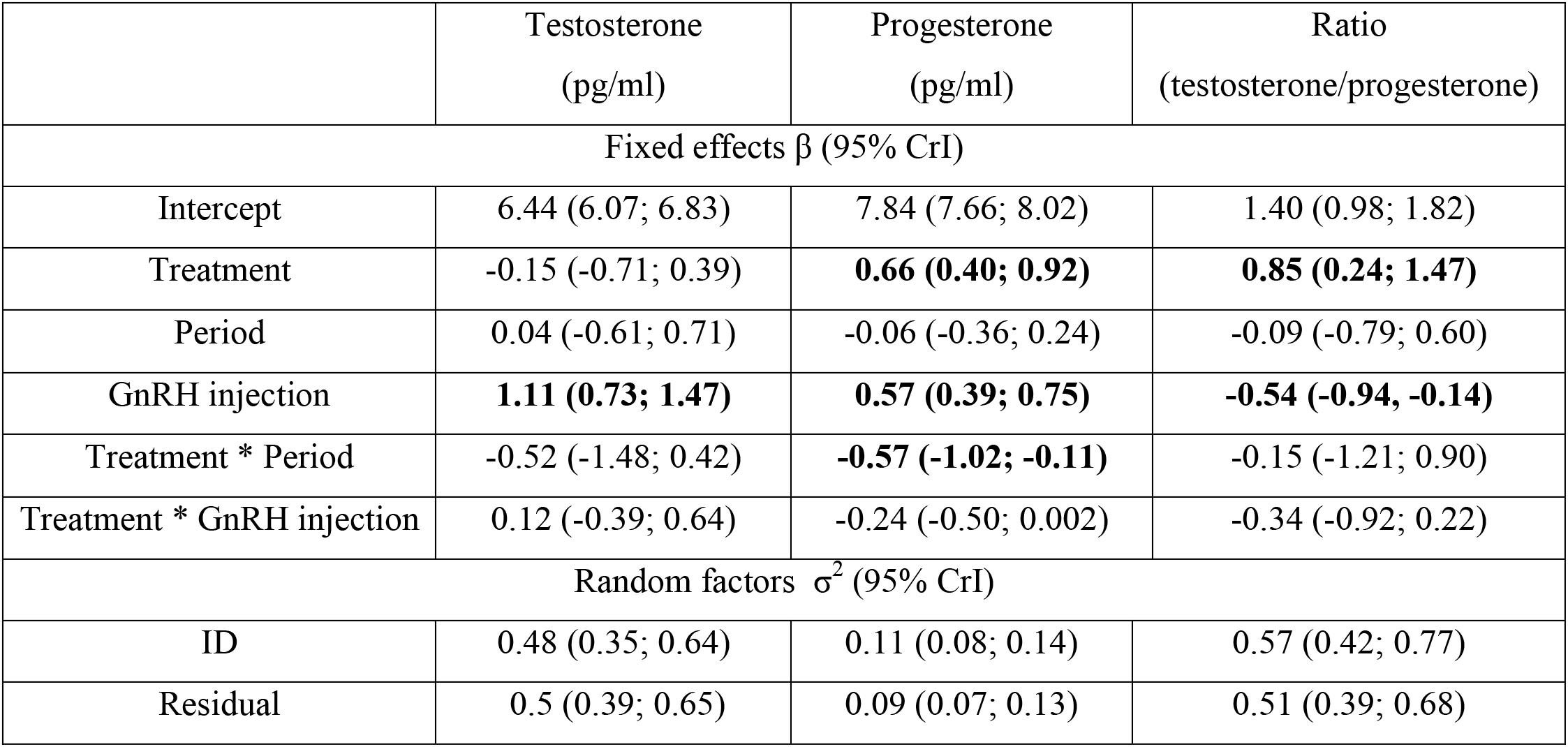
Results from a linear mixed effect model estimating fixed and random effects to explain variation in rufous hornero hormone levels. Treatment (control vs. STI), period (mating vs. parental care period), GnRH injection (pre vs. post GnRH injection), capture and handling time were fitted as fixed factors. The interactions treatment:period and treatment:injection were also included. We present fixed (β) and random (σ^2)^ parameters with their 95% credible intervals (CrI) in brackets. A statistically meaningful effect of a fixed factor can be assumed if zero is not included within the 95% CrI or if mean difference between compared estimates is higher than 0.95, and are presented in bold font.

|  | Testosterone<br>(pg/ml) | Progesterone<br>(pg/ml) | Ratio<br>(testosterone/progesterone) |
| --- | --- | --- | --- |
| Fixed effects $\beta$ (95% CrI) | | | |
| Intercept | 6.44 (6.07; 6.83) | 7.84 (7.66; 8.02) | 1.40 (0.98; 1.82) |
| Treatment | -0.15 (-0.71; 0.39) | **0.66 (0.40; 0.92)** | **0.85 (0.24; 1.47)** |
| Period | 0.04 (-0.61; 0.71) | -0.06 (-0.36; 0.24) | -0.09 (-0.79; 0.60) |
| GnRH injection | **1.11 (0.73; 1.47)** | **0.57 (0.39; 0.75)** | **-0.54 (-0.94, -0.14)** |
| Treatment * Period | -0.52 (-1.48; 0.42) | **-0.57 (-1.02; -0.11)** | -0.15 (-1.21; 0.90) |
| Treatment * GnRH injection | 0.12 (-0.39; 0.64) | -0.24 (-0.50; 0.002) | -0.34 (-0.92; 0.22) |
| Random factors $\sigma^2$ (95% CrI) | | | |
| ID | 0.48 (0.35; 0.64) | 0.11 (0.08; 0.14) | 0.57 (0.42; 0.77) |
| Residual | 0.5 (0.39; 0.65) | 0.09 (0.07; 0.13) | 0.51 (0.39; 0.68) |

#### Aggression in relation to hormones

We performed a linear model for each hormone and the progesterone-to-testosterone ratio using log-transformed variables. As explanatory variables, in all cases, we used the breeding stage (mating vs. parental care period) and the normalized ‘number of flights over the decoy’, which we have shown to be the most reliable proxy for territorial aggression in this species (Adreani et al. unpublished). The ‘number of flights over the decoy’ was normalized between 0 and 1 for each period in order to have comparable aggression gradients across periods, optimizing the model fit. Because the relationship between aggressive behavior and hormone concentration could differ between periods, we included this interaction. We also included handling time as a covariate in the models for progesterone and the ratio. For details on these models see Table 2.

**Table 2.**
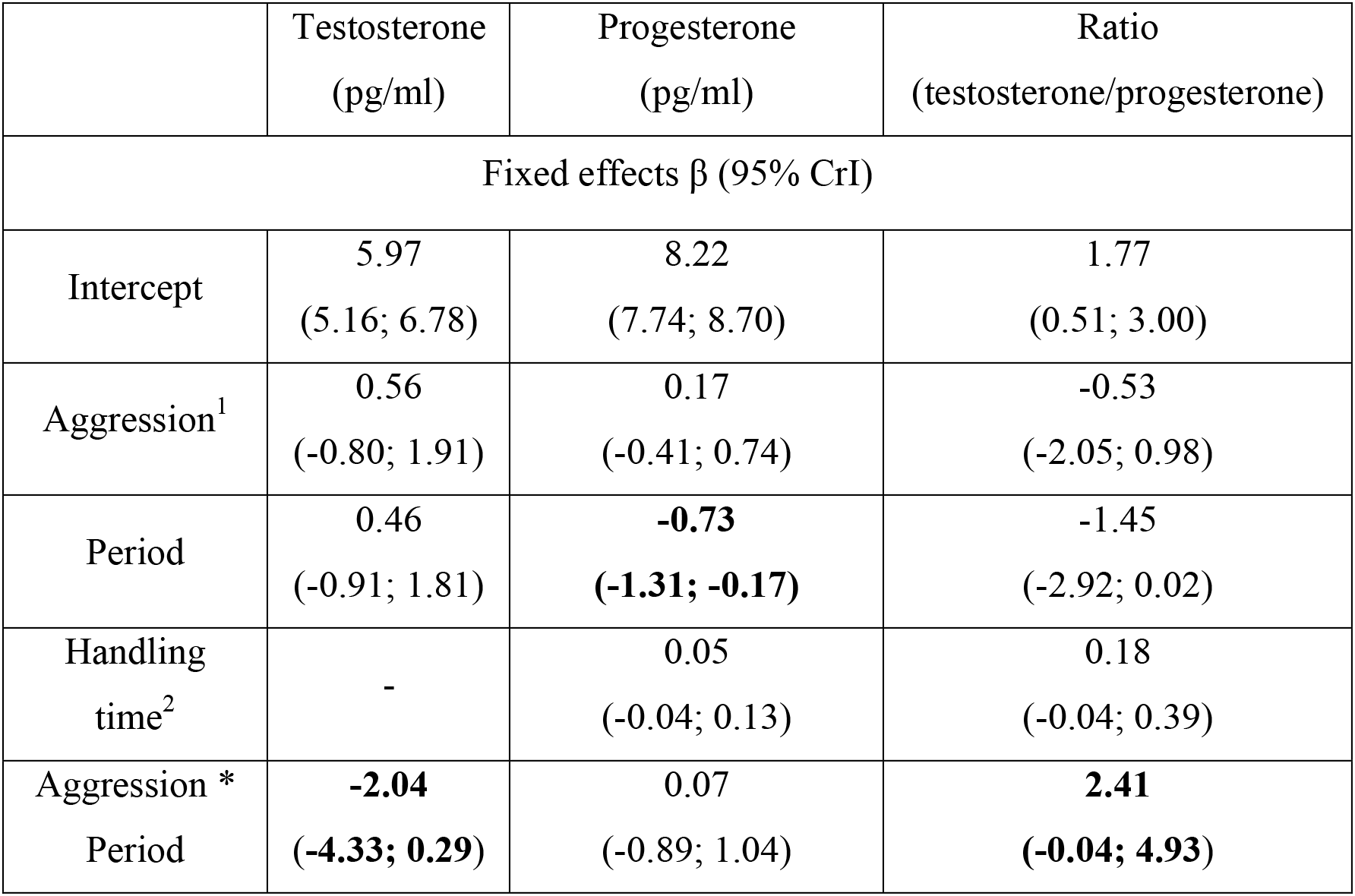
Results from a linear model estimating fixed effects to explain variation in rufous hornero hormone levels. Aggression (“flights over the decoy”), period (mating vs. parental care period) and handling time (not for testosterone) were fitted as fixed factors. The interaction between aggression and period was also included. We present fixed (β) parameters with their 95% credible intervals (CrI) in parentheses. A statistically meaningful effect of a fixed factor can be assumed if zero is not included within the 95% CrI or if mean difference between compared estimates is higher than 0.95, and are presented in bold font.

## Results

### Effect of STI on hormonal levels

STI and control males did not differ in post-capture testosterone levels during the mating period (Fig. 1A; Table 1), but STI birds had higher progesterone levels after capture (Fig. 1B; Table 1). During parental care, STI birds had lower testosterone levels than controls (Fig. 1A; Table 1), but progesterone levels did not differ between the two groups (Fig. 1B; Table 1). Finally, the progesterone-to-testosterone ratio was higher in STI birds than in controls during both periods (Fig. 1C; Table 1). The progesterone-to-testosterone ratio of controls and STIs did not differ between periods (Fig. 1C; Table 1).

**Figure 1.**
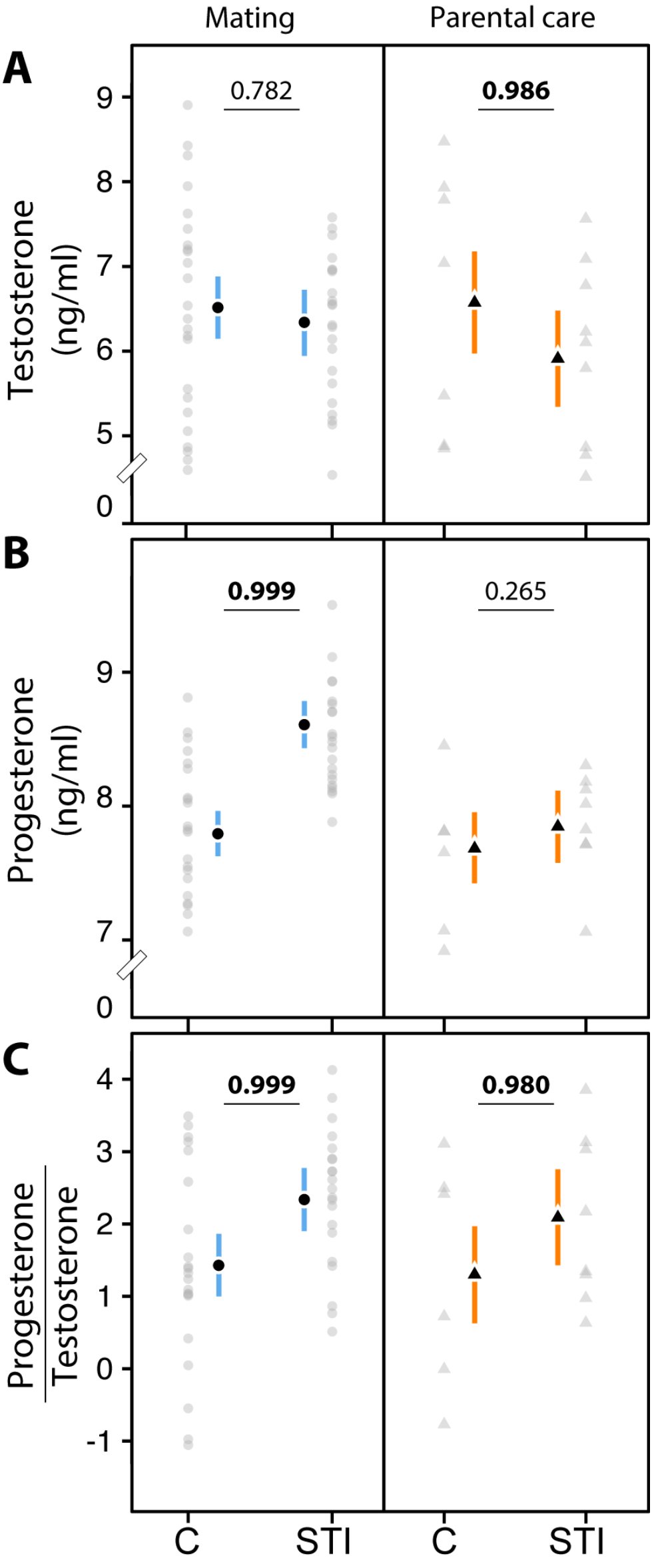
Log-transformed levels of (a) testosterone and (b) progesterone, and (c) the ratio of progesterone to testosterone control (C) and challenged (STI) male horneros. Grey dots and triangles represent raw data; black dots and triangles the predicted estimates of the models and the vertical bars the 95% CrI. Numbers represent the posterior probability of a mean difference between C and STI birds, and in bold are the numbers with a posterior probability higher than 0.95.

In both periods control and STI birds increased testosterone and progesterone concentrations after injection of GnRH (Suppl. Fig. 1; Table 1).

### Relationship between aggression and hormones

The number of flights over the decoy was not related to post-capture testosterone, progesterone, or the ratio of these hormones during the mating period (Figure 2 A, B, C; Table 2). During parental care, the number of flights over the decoy was unrelated to post-capture progesterone (Figure 2 B; Table 2), but the number of flights and post-capture testosterone were negatively related, with more aggressive birds expressing lower levels of testosterone (Figure 2 A; Table 2). This negative relationship also drove a positive relationship between the ratio of progesterone to testosterone and the number of flights over the decoy (Figure 2 C; Table 2). This differential effect of testosterone and the progesterone to testosterone ratio during mating and parental care was reflected by the effect of the interaction term between aggression and period (Figure 2 D, F; Table 2).

**Figure 2.**
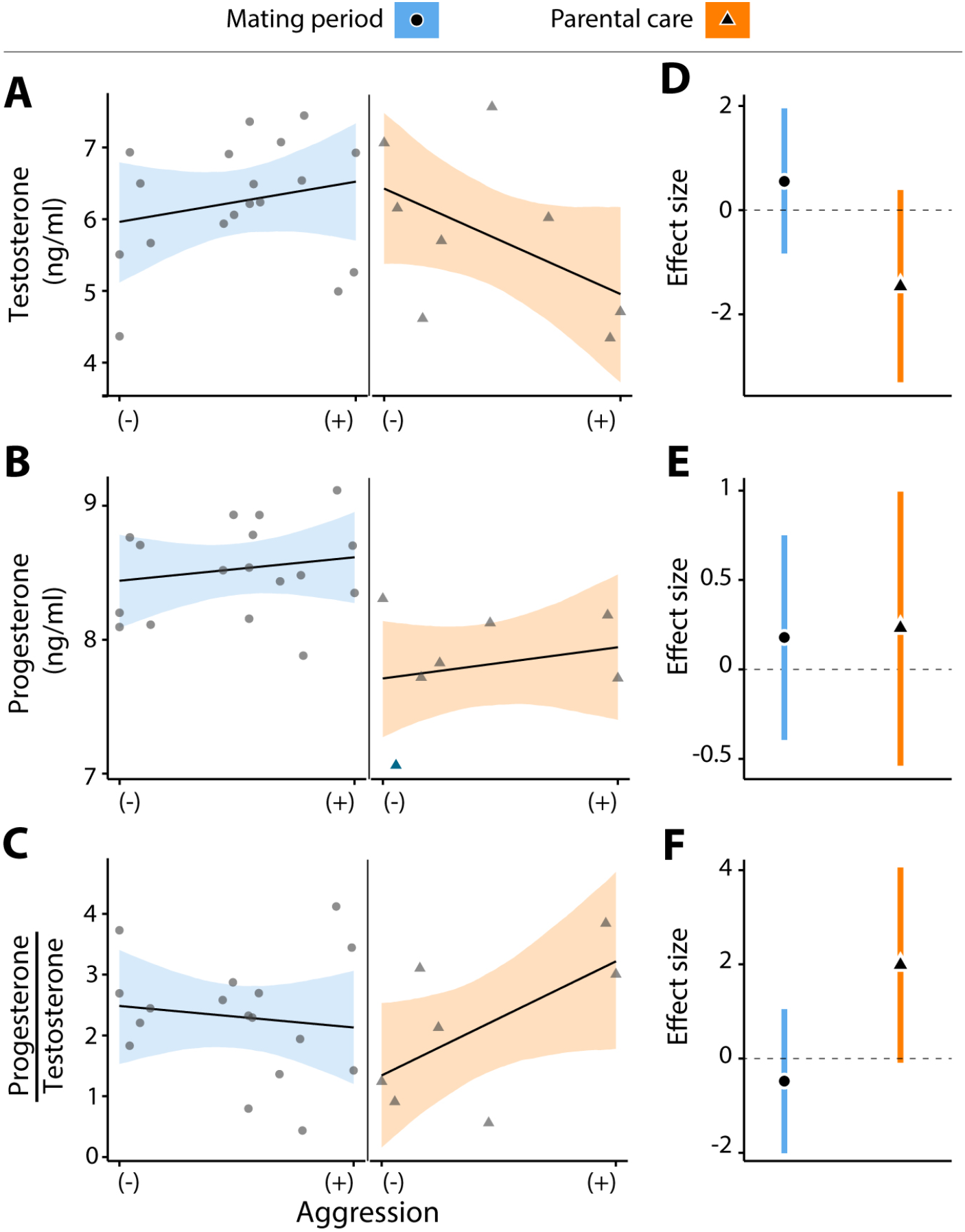
Relationship between the number of flights over decoy (a proxy for aggressiveness) and testosterone (A), progesterone (B) and their ratio (C) during mating and parental care periods. Black shapes represent mean estimates of the models and ribbons represent 95% CrI. Grey shapes represent raw data. Hormonal and ratio values are log-transformed. Also depicted in (D), (E) and (F), the respective effect sizes (i.e., slopes) for the models. Black shapes represent the mean slope values and the vertical bars the 95% CrI.

## Discussion

In free-living male horneros, we did not find support for the ‘challenge’ or the ‘progesterone challenge’ hypothesis. However, simulated territorial intrusions induced a similar change in the progesterone to testosterone ratios during the mating and the parental care period by differentially affecting both hormones in each period.

In male horneros, testosterone did not increase as a consequence of male-male aggression as predicted by the ‘challenge hypothesis’ (Wingfield et al., 1990). The subsequent rise in testosterone after injection of GnRH indicated that they had the physiological capacity to increase testosterone (Supplementary Figure 1). Hence, our study adds to the growing evidence that testosterone levels do not rise during male-male challenges in most species of biparental birds (reviewed by Goymann 2009 and Goymann *et al.* 2015). Also, the level of aggression was unrelated to testosterone during the mating phase, but during parental care more aggressive males paradoxically had lower testosterone. Similar findings have been reported in a study by Apfelbeck & Goymann (2011) and such pattern were attributed to a potentially higher metabolism during STIs. Given the low sample size during parental care, the correlation we found should be considered with caution.

Only few experimental studies have investigated the link between progesterone and aggression in males (e.g. Erpino & Chappelle 1971; Fraile *et al.* 1987, 1988;Schneider *et al.* 2003; Fuxjager *et al.* 2009) and none of them did in free-living animals. Further, if aggressive behavior *per se* affects the progesterone pathway remained unknown. In male horneros STI birds had higher progesterone levels than controls during the mating period but not during the parental care period. In male house mice (*Mus musculus*) (Schneider et al., 2003) and male tree lizards (*Urosaurus ornatus*) more aggressive individuals also have higher levels of progesterone (Weiss and Moore, 2004) and male California mice (*Peromyscus californicus*) increase progesterone after winning aggressive encounters (Fuxjager et al., 2009). These data support a role of progesterone in modulating male aggressive behavior (but see Erpino and Chappelle, 1971; Fraile et al., 1987), and our results are the first to show that social interactions can modulate progesterone in males. In addition, our findings suggest a sexually distinct modulation of progesterone by aggressive behavior, because in female birds aggressive behavior either decreases (Goymann et al., 2008) or does not affect progesterone (Elekonich and Wingfield, 2000). This could possibly be due to the different roles that progesterone plays in each sex (Brinton et al., 2008). Thus, it is probable that the ‘progesterone challenge hypothesis’ that was proposed for females (Davis and Marler, 2003) cannot be extended to males of other vertebrate species. Interestingly female horneros, as males, are also aggressive year-round (Fraga, 1980). Given that progesterone and testosterone can also play a role in agonistic behaviors in females (Cooper and Crews, 1987), it could be that our findings could be extended to female horneros. However, only three studies have experimentally addressed the effect of aggression on progesterone in females (Davis and Marler, 2003; Elekonich and Wingfield, 2000; Goymann et al., 2008), and ours represent the first study in males and across life-cycle stages, thus further studies are necessary to better understand how aggression interacts with progesterone across vertebrates.

### GnRH and progesterone

GnRH induced a progesterone response in control and challenged birds during both periods. This first report of such effect of GnRH is relevant in itself besides the fact that all the horneros had the physiological capacity to increase progesterone. GnRH injections have been used as treatment to increase testosterone within an individual’s range (e.g. Goymann and Flores Dávila, 2017), thus in the future when interpreting its effect on behavior is important to consider that also progesterone might be affected too.

### Progesterone to testosterone ratio and aggression

Given the evidence of joint actions of progesterone and testosterone altering sexual behavior in males (Erickson et al., 1967; Erpino and Chappelle, 1971; Fraile et al., 1988, 1987; Fuxjager et al., 2009; Schneider et al., 2003), behavior could also feedback on both hormones, allowing for their interaction. The progesterone-to-testosterone ratio reflects the balance between both hormones at an individual level. Although aggression affected each hormone differentially across periods and contrary to our predictions, it induced a similar increase in the ratio of progesterone to testosterone in both periods (Fig. 1C). In lizards, progesterone potentiates the effects of testosterone by altering the expression of androgen and progesterone receptors (Godwin and Crews, 1997). Similar mechanisms could potentially be operating in birds: behavior could trigger the differential release of testosterone and progesterone during different breeding sub-stages to orchestrate the physiology of a bird when challenged by a rival. Given that both hormones can interact, our results open the exiting possibility that an effective physiological response resides in the relation between both hormones rather than in the concentration of each independently.

### Conclusion

Overall, we show for the first time in males of a free-living vertebrate, that simulated territorial intrusions affected testosterone and progesterone differentially depending on whether the animals were in the mating or in the parental care period. This differential change in hormone levels resulted in a similar change of the ratio between progesterone and testosterone, independent of the life-cycle stage. Our findings suggest that the physiological response to aggressive interactions might not be limited to one hormone, but may lead to an orchestrated response that differs between life-cycle stages. New studies on multiple hormones are needed to further investigate this exciting perspective.

## Ethics

The Ethics Committee of Animal Experimentation (CUEA) of the Universidad de la República de Uruguay approved all experimental procedures. Protocol number 186, file 2400-11000090-16.

## Data accessibility

The dataset supporting this article will be available at Mendeley Data.

## Authors contributions

NMA and LM share the first and last authorship. NMA and LM conceived the study, designed the experiment, collected and analyzed the data. WG advised regarding the experimental design and interpretation of the results. NMA wrote the first draft of the manuscript, which was then revised by all authors. All authors approved the final version of the manuscript.

## Acknowledgements

We are grateful to Manfred Gahr and Michaela Hau for their support, Enzo Cavalli and Ernesto Guedes for their assistance in the field, and Monika Trappschuh for conducting the hormone analyses. We thank the Ethology Lab at the Universidad de la República in Uruguay, and Bettina Tassino for her support. We thank the ‘INIA Las Brujas’ staff in Canelones, Juan Carlos Reboreda and LEyCA lab from the University of Buenos Aires. We also thank Pablo Tubaro from the Museo Argentino de Ciencias Naturales ‘Bernardino Rivadavia’ (MACN) for providing us with the mounted hornero, and Michaela Hau for constructive feedback on earlier versions of the manuscript.

---

**Supplementary figure 1.**
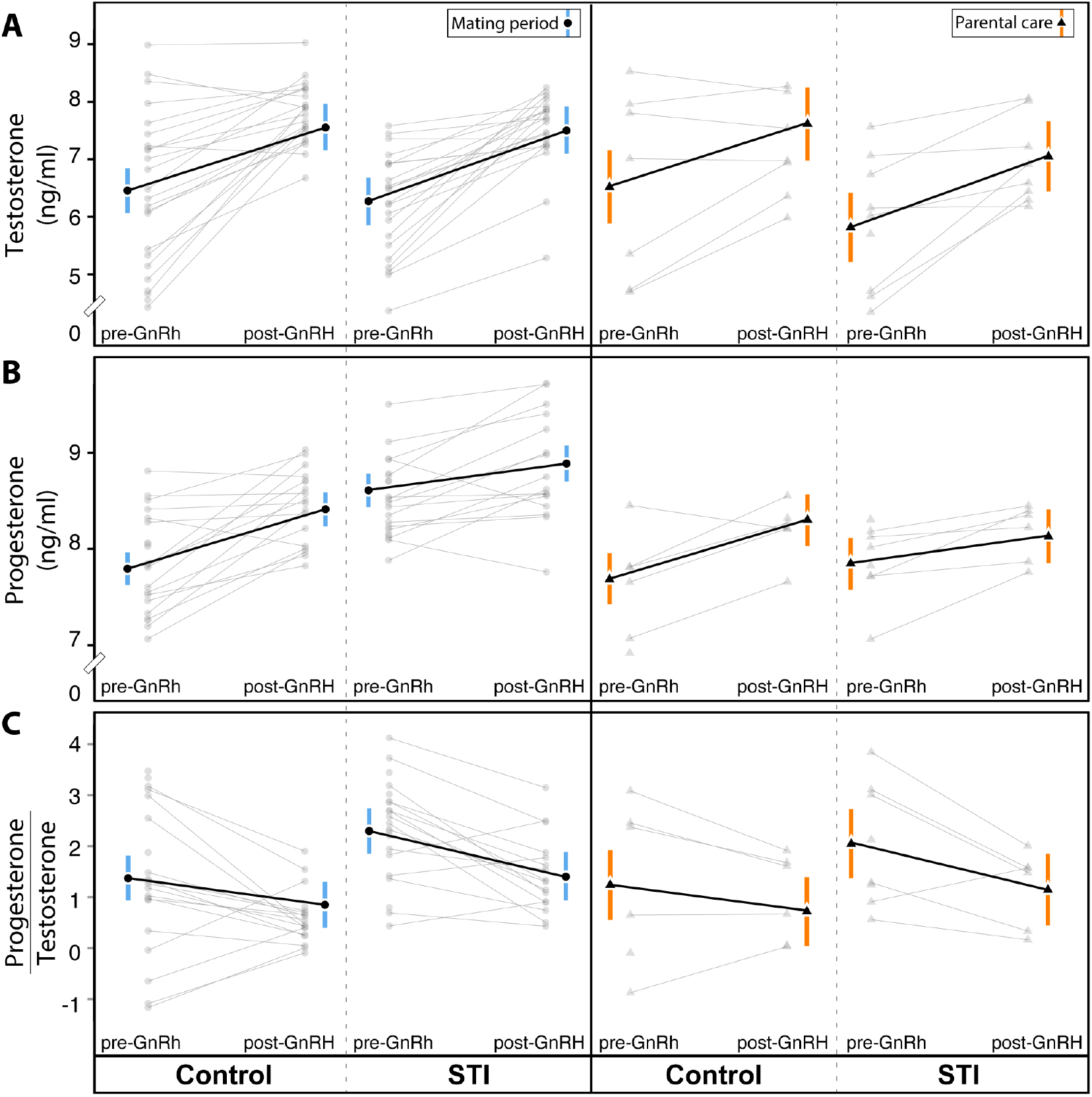
Pre and post GnRH induced (a) testosterone and (b) progesterone levels and (c) progesterone/testosterone ratio of ‘control’ and ‘STI’ males during mating and parental care period. Black shapes represent mean estimates of the models and vertical bars represent 95% CrI. Grey shapes represent raw data, black lines the estimated change and grey lines the change in concentration of each individual after the injection. Hormonal and ratio values are log-transformed.

**Table S1.** Results from a linear model estimating fixed effects to explain variation in rufous hornero body condition. Treatment (control vs. STI) and period (mating vs. parental care period) were fitted as fixed factors together with the interaction between them. We present fixed (β) parameters with their 95% credible intervals (CrI) in parentheses. A statistically meaningful effect of a fixed factor can be assumed if zero is not included within the 95% CrI or if mean difference between compared estimates is higher than 0.95.

|  | Mass (gr) | Tarsus (mm) | Wing (mm) |
| --- | --- | --- | --- |
| Fixed effects $\beta$ (95% CrI) | | | |
| Intercept | 58.20<br>(49.93; 66.44) | 34.04<br>(31.45; 36.61) | 10.72<br>(9.89; 11.54) |
| Treatment | -0.94<br>(-12.58; 10.66) | -0.74<br>(-4.33; 2.86) | -0.33<br>(-1.51; 0.82) |
| Period | -0.16<br>(-2.69; 2.38) | -0.08<br>(-0.86; 0.72) | -0.05<br>(-0.29; 0.21) |
| Treatment *<br>Period | 0.34<br>(-3.19; 3.87) | 0.27<br>(-0.83; 1.38) | 0.09<br>(-0.27; 0.45) |

**Table S2.** Results from a linear model estimating fixed effects to explain variation in experimental groups’ sampling and capture time. Treatment (control vs. STI) and period (mating vs. parental care period) were fitted as fixed factors together with the interaction between them. We present fixed (β) parameters with their 95% credible intervals (CrI) in parentheses. A statistically meaningful effect of a fixed factor can be assumed if zero is not included within the 95% CrI or if mean difference between compared estimates is higher than 0.95.

|  | Sampling Time (s) | Capture Time (min) |
| --- | --- | --- |
| Fixed effects $\beta$(95%CrI) | | |
| Intercept | 337.97<br>(278.24; 396.56) | 4.48<br>(2.78; 6.20) |
| Treatment | -53.01<br>(-140.75; 35.07) | 1.63<br>(-0.99; 4.22) |
| Period | 16.84<br>(-109.06; 137.38) | 0.10<br>(-3.47; 3.72) |
| Treatment * Period | 49.01<br>(-120.64; 216.83) | 0.26<br>(-4.85; 5.30) |

